# Graphia: A platform for the graph-based visualisation and analysis of complex data

**DOI:** 10.1101/2020.09.02.279349

**Authors:** Tom C. Freeman, Sebastian Horsewell, Anirudh Patir, Josh Harling-Lee, Tim Regan, Barbara B. Shih, James Prendergast, David A. Hume, Tim Angus

## Abstract

Quantitative and qualitative data derived from the analysis of genomes, genes, proteins or metabolites from tissue or cells are currently generated in huge volumes during biomedical research. Graphia is an open-source platform created for the graph-based analysis of such complex data, e.g. transcriptomics, proteomics, genomics data. The software imports data already defined as a network or a similarity matrix and is designed to rapidly visualise very large graphs in 2D or 3D space, providing a wide range of functionality for graph exploration. An extensive range of analysis algorithms, routines for graph transformation, and options for the visualisation of node and edge attributes are also available. Graphia’s core is extensible through the deployment of plugins, supporting rapid development of additional computational analyses and features necessary for a given analysis task or data source. A plugin for correlation network analysis is distributed with the core application, to support the generation of correlation graphs from any tabular matrix of continuous or discrete values. This provides a powerful analysis solution for the interpretation of high-dimensional data from many sources. Several use cases of Graphia are described, to showcase its wide range of applications. Graphia runs on all major desktop operating systems and is freely available to download from https://graphia.app/.

## Introduction

The study of interactions between entities is a cornerstone of modern analytics. In biology, efforts to map the ‘interactome’, all the interactions between the components of a biological system, have been underway for some time, generated by a number of complementary approaches^1,2^. Networks of biological data may also be used to chart diverse phenomena such as the spread of disease, the interactions between drugs and their targets, and the evolutionary relationships between species. Many data from other sectors are also inherently graph-based in structure. For example, interactions on social media platforms, customer-client relationships, communication and transport systems, computer networks and many other real-world systems, can all be considered as the edges and nodes of a graph. Matrices of numerical data that do not inherently possess a network structure can also be analysed using graph-based approaches. Wherever it is possible to calculate the distance between entities, a graph can be constructed using high confidence measures to define the edges between entities, represented by nodes. In biology, such an approach is already widely used to analyse high dimensional data, in particular to construct and analyse gene coexpression networks^3,4^, but the approach is applicable to any numerical or categorical data from any source.

Given the explosion in the availability of data in recent years and the potential to visualise and analyse it using graph-based approaches, a variety of software tools to support these activities have been developed. In biology, Cytoscape^5^ is the most widely used software for performing graph analytics. It has a large user base and supports many ‘apps’ (plugins) created by the community for the performance of specific graph-based analysis tasks. Other network visualisation and analysis tools include BioLayout^4,6^ Gephi^7^, Graphviz^8^, Pajek^9^, yEd (yFiles, Tübingen, Germany), Social Network Visualiser^10^ and NodeXL^11^. There are also a range of web-based software tools exclusively designed to visualise portions of data, often from a designated database, such as String^12^, GeneMania^13^ and Neo4J Bloom^14^. Some of these tools are focused on supporting a particular community, whilst others possess functionality tailored towards specific tasks or data types and include a mix of open-source projects and commercial tools. Others provide open-source code repositories for graph visualisation and analysis algorithms,^15^ or share repositories of graph data^16–18^. For a more comprehensive review of network analysis tools and resources, see^19^.

Despite the availability of a wide range of downloadable applications, web-resources and code libraries to support graph-based analyses, there is a pressing need for easy-to-use software that supports the rapid visualisation and analysis of relatively massive networks. To address this need, we developed Graphia - a general purpose graph analysis tool that supports the integration, visualisation, analysis, and interpretation of a wide variety of data types. Here we provide an overview of Graphia’s core functionality for the analysis of graphs and describe a number of case studies in which it is applied to solve problems associated with the analysis of data derived from the biological sciences.

## METHODS AND RESULTS

### Design Criteria

The following features were considered core to the design of Graphia:

- ***Data and operating system agnostic.*** Import data from any source saved in standard file formats. The software should run on all major desktop operating systems and modern hardware configurations.
- ***Fast and scalable.*** Support the rapid loading of data, fast computation of graph layout and analysis algorithms, high quality data visualisations. Deliver smooth and responsive graphical rendering of millions of data points (node/edges) on standard desktop hardware.
- ***Dynamic rendering.*** Visualise in real time changes to the graph structure associated with alterations in input parameters or additional data.
- ***3D graph visualisation.*** Provide a navigable and immersive environment in which to explore and interpret large and complex graph topologies.
- ***Correlation graphs as an essential function.*** Rapidly convert any numerical or categorical data table into a correlation graph, supporting pattern finding and data mining.
- ***Attribute handling and visualisation***. Visualise attributes (metadata) associated with nodes and edges using colour, size and text to distinguish between attribute values.
- ***Advanced analysis capabilities.*** Support a wide range of analytical algorithms and approaches that empower a user to explore, query and interpret data.
- ***Extensible.*** Provide extensible architecture through use of a plugin system to allow the core to be extended or adapted for specific application areas or data types.
- ***User Interface.*** Provide a simple and intuitive user interface (UI) that is easy to navigate, featuring a graph display area supplemented with a table listing selected nodes and associated attributes and data values. The UI should provide easy access to menus providing functionality and display active transformations and visualisations.

### Code Architecture

Graphia is written in C++17 and is built upon Qt version 5, the cross-platform widget toolkit. For graphics, the industry standard OpenGL is used. The minimum driver support required is version 3.3 core profile, but more modern extensions will be used if they are available. Various open source libraries are employed, mostly for loading external data formats. These libraries and their associated licenses are enumerated in the About dialog of the application, accessible from the Help menu.

Graphia is architected so that loading, and data type-specific user interfaces are confined to plugins. These are independent modules to the core application and can be removed or added without affecting any base functionality.

At the highest level the code is organised hierarchically into four separate directories:

- app - the core application code
- plugins - the existing bundled plugins
- shared - code used by both the core and plugins (this includes interface headers)
- thirdparty - any library code not authored locally

These are further divided into subdirectories dealing with specific areas of functionality. Graphia has been developed using standard object-orientated best practices. Continuous integration is employed to prevent portability build regressions, using recent versions of the compilers GCC, clang and MSVC. In addition, static analysis tools such as clang-tidy and cppcheck are used to identify potential problems early. CMake is used as a build system, and is set up for Linux, Windows and MacOS compilation.

### Implementation

#### Data Import

Graphia has been designed to import data encoded in a variety of standard and non-standard file formats. These include standard graph-based file formats such as BioPAX OWL ontology (.owl), JSON graph (.json), GraphML (.graphml) and Graph Modelling Language (.gml), MATLAB data file (.mat) formats, and non-standard but simple formats such as pairwise graph formats (.txt, .layout), adjacency matrix (.csv, .tsv) and numerical data prepared for correlation analyses (.csv, .xls). Using these file formats, a wide variety of data may be imported into Graphia, not only in terms of defining the nodes and edges of a graph, but user-defined attributes or metadata.

#### Fast and Scalable

Existing graph visualisation tools either fail to render very large graphs effectively, or the ability to interact with a graph once rendered is limited and slow. Therefore, all aspects of Graphia’s functionality have been engineered to run quickly. Graphia can render graphs millions of data points on relatively commonplace hardware, where interaction with them is fast and fluid. This has been achieved through the use of optimal coding practices and parallelisation of computationally intensive analysis routines, e.g. calculation of correlation matrices, graph layout, clustering.

#### Dynamic Graph Layout and Rendering

Graph layout is an iterative process. Many programs only display the results of a layout algorithm after it has run a defined number of iterations. With Graphia, the layout is shown live, such that graphs ‘unfold’ in real time. However, the true power of dynamic graphs is realised when a transformation operation is performed or following the addition of new data. These changes are immediately reflected in the appearance of the graph. As graphs change dynamically, there is a need to identify the graph components and map how they interact when construction parameters are adjusted. The ability to quickly identify and move between components is a unique feature of Graphia. Components are rendered in a concentric pattern, arranged large-to-small. Smaller components may be filtered away using a transform (see below).

Most existing network analysis tools render graphs in 2D. The layout algorithm implemented in Graphia is innovative in that it applies current force-directed layout techniques, but in a dynamic setting. Graphia renders graphs in 3D or 2D, making use of modern graphics hardware to display extremely large graphs efficiently with options for node/edge shading, relative node sizing and spacing (Figure 1A-F). When graphs are relatively small or there is a need to share images by a conventional medium, i.e. a document, 2D graph visualisations have advantages. However, 2D visualisations are limiting when there is a need to display and interact with large graphs with complex topologies.

**Figure 1.**
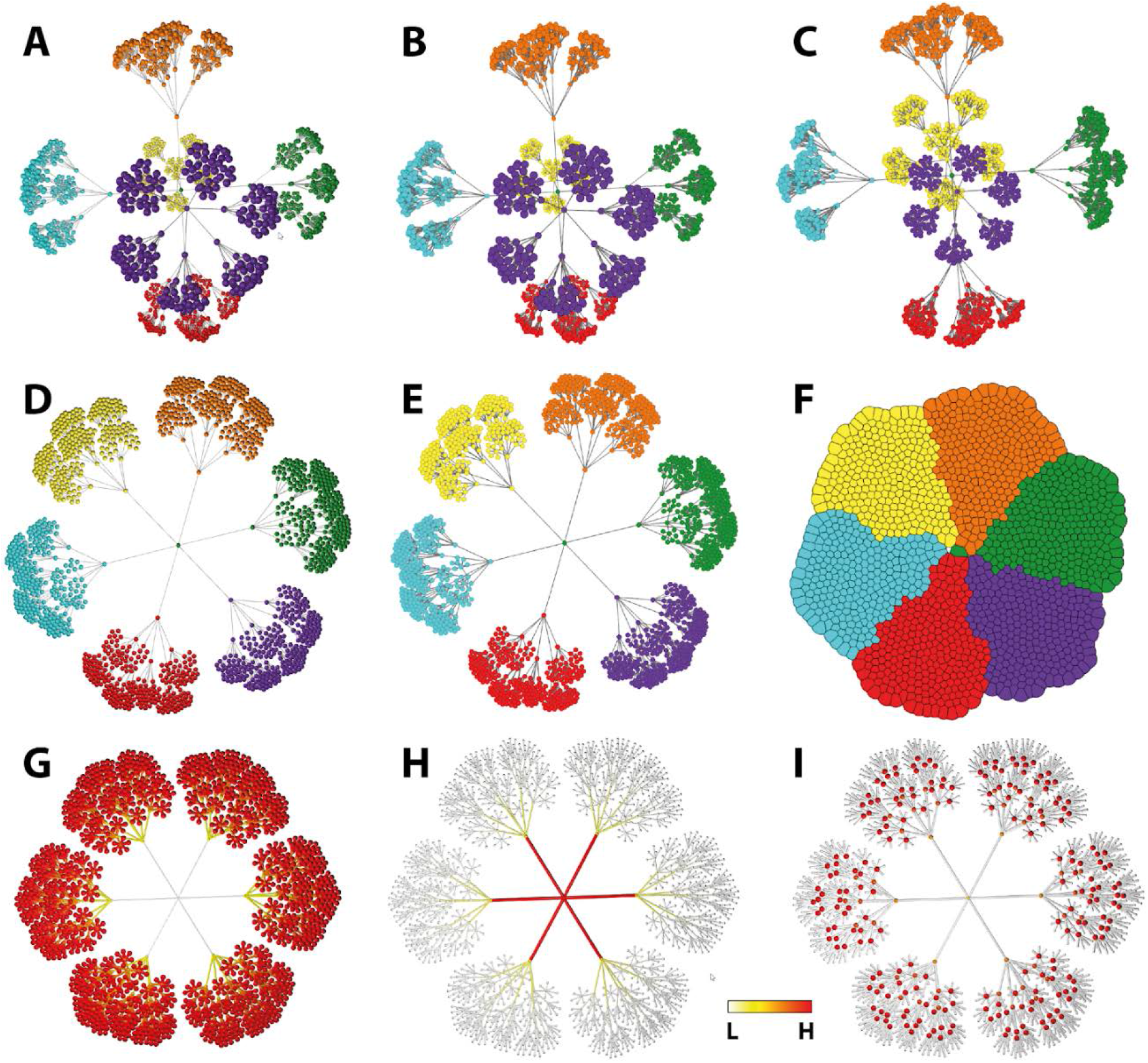
Different graph visualisation options. **(A)** 3D perspective view, smooth shading (the default), with visualisation of node categorical attribute (MCL cluster). **(B)** 3D perspective view, flat shading. **(C)** 3D orthographic view, flat shading (no perception of distance - all nodes same size, unless sized by attribute value). **(D)** 2D view, smooth shading. **(E)** 2D view, flat shading. **(F)** compressed 2D layout, flat shading, showing node overlap view. **(G)** Visualisation of Betweeness centrality values, **(H)** Eccentricity values, **(I)** PageRank values. G-I are continuous (numerical) attributes, so a colour spectrum and size gradient is used for node display (2D, smooth shading). Betweenness and eccentricity are calculated for both nodes and edges, therefore visual encoding is applied to both.

#### Attribute-to-Visual Mapping

Attributes are data values associated with nodes/edges. These can be user-defined or calculated by Graphia. For example, a node representing a person may be associated with knowledge of their gender, occupation, socioeconomic class, ethnicity, etc. (categorical attributes), as well as their height, age, weight, years in employment (continuous value attributes). Colour can be used to represent categorical attributes, with nodes sharing the same attribute being assigned the same colour. In the case of continuous value attributes, colour and size can be used to represent the value according to a spectrum, e.g. from small white nodes to big red nodes to represent low and high values, respectively. Both types of attribute may also be calculated from the graph itself, e.g. the assignment of nodes to clusters or calculation of node degree, PageRank values, etc., respectively (Figure 1G-I). Visualising attribute values may help explain graph structure, for instance an area of a graph might be visibly associated with nodes of a given attribute. Attributes may also be used to analyse the statistical associations with graph topology, for example, graph clusters may be analysed for the enrichment of nodes with a given attribute. Attribute values may also be specific to only a single element, e.g. a unique node name.

The ‘Add Visualisation’ menu (Figure 2.2a) provides a highly flexible interface to generate and modify attribute-to-visual mappings. Enrichment analysis of attributes associated with a graph is available under the ‘Tools’ menu.

**Figure 2.**
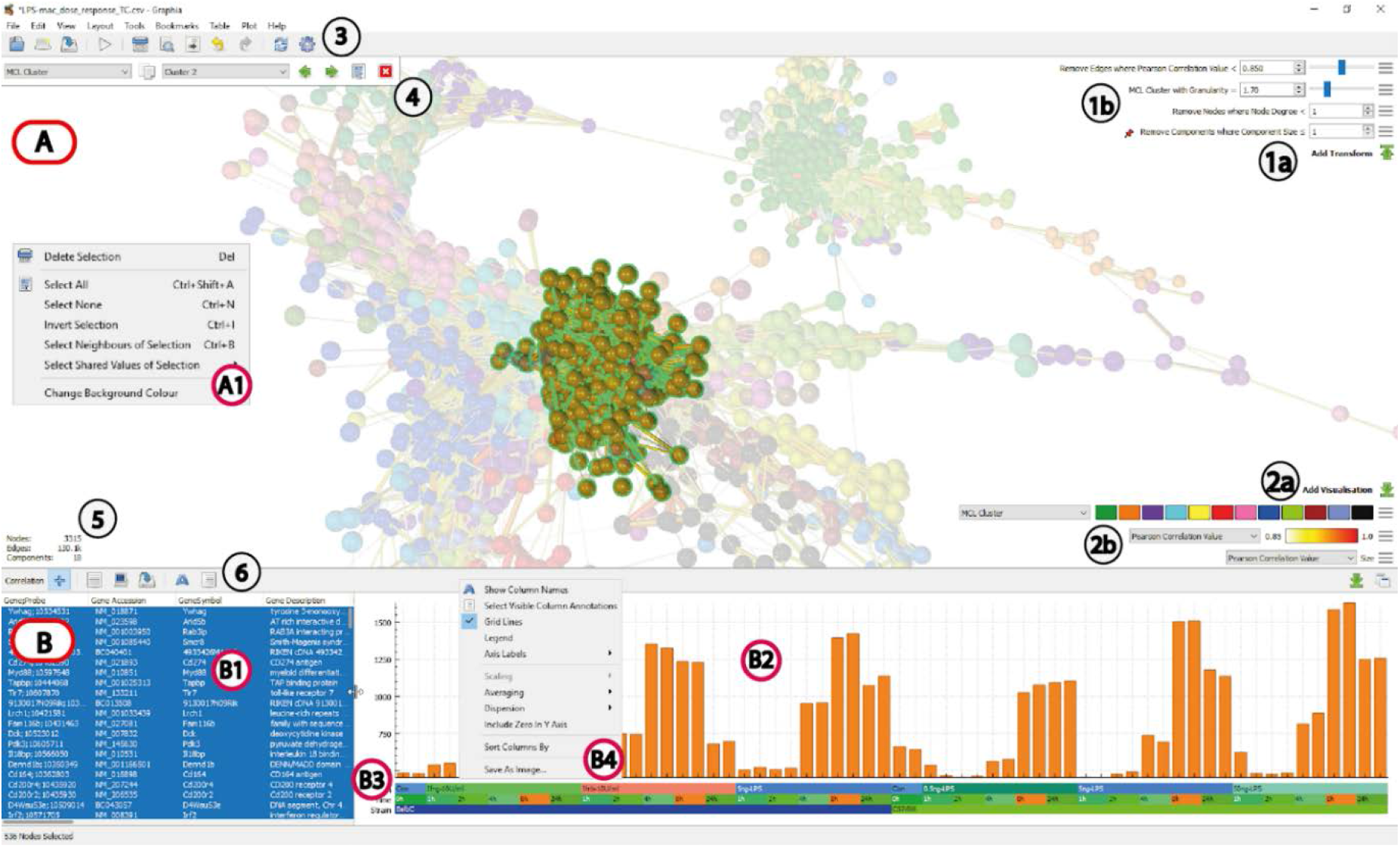
Graphia user interface displaying a correlation graph. **(A)** Graph display area, showing correlation graph with a cluster selected (unselected nodes faded). **(A1)** Display context menu options (right click). **(B)** Node (row) attribute display area. **(B1)** Table of selected nodes and their attributes. **(B2)** Data plot area showing average profile of selected nodes. **(B3)** Visualisation of column annotations. **(B4)** Data plot context menu options for changing plot (right click). **(1a)** Add Transform button, **(1b)** Active transforms, **(2a)** Add visualisation button, **(2b)** Active visualisations, **(3)** General toolbar, **(4)** Attribute parameter selection, **(5)** Display of graph metrics (number of nodes, edges, components), **(6)** Plot/table function toolbar.

#### Graph Transformation and Analysis

A transform is defined as a process by which a graph’s structure or data is altered. Options for transforming a graph are available through the ‘Add Transform’ button (Figure 2.1a). Clicking this button opens a dialog in which a transform may be selected, and its parameters configured. Structural changes may be brought about through the application of algorithms that for instance remove ‘branches’ or ‘leaves’ from a graph, or contract edges based on an attribute value. Similarly, filters may be applied that remove edges below a certain weight or allow a user to remove nodes where their node degree is greater or lesser than a certain value. One further type transformation available is the ability to reduce a graph’s quantity of edges. In some graphs the ratio of edges to nodes can be very high and can obscure higher order topologies, as well as using additional resources to render them. Graphia incorporates the k-NN algorithm^20^, which culls all but the top *k* edges, according to the value of a user nominated attribute. %-NN is a variation of this that instead retains a defined percentage of the original edges. Finally, there is an algorithm that simply removes a homogenous random sample of edges, for the case where no edge attribute is available. Changes to graph structure brought about by the application of any of the above algorithms or filters are immediately visible through the dynamic layout feature.

Graphia automatically generates node-level analyses of any graph, calculating the node degree, and multiplicity of individual nodes/edges. Where a graph has been generated from a numerical matrix, it will also automatically calculate the maximum, minimum mean and variance of the data series represented by a node. Topological analyses can be achieved through clustering and Graphia incorporates the MCL^21^ and Louvain^22^ clustering algorithms, where the granularity of clustering can be adjusted after their initial calculation. The MCL algorithm is optimal for highly structured graphs where the ratio of edges to nodes may be high, e.g. correlation graphs. The Louvain algorithm is favoured when working with sparse graphs where the ratio of edges to nodes may be low, e.g. after the application of an edge reduction algorithm such as k-NN. After the assignment of nodes into clusters, Graphia can then be used to perform an enrichment analysis to test the hypothesis that clusters may be enriched with nodes possessing specific a given attribute. The option to perform enrichment analyses is found under the Tools menu.

#### Plugins

To allow the functionality of the core application to be extended to perform a specific task or work with data from a specific resource, Graphia uses a plugin architecture. A plugin may include additional algorithms and network analyses methods or be designed to import/export data from a cloud-based resource or database or guide a user through a specific analysis routine. In the case of Graphia’s ‘correlation plugin’, this is specifically designed for the import of numerical matrices and their conversion into a correlation graph, where nodes represent data series and edges represent high confidence relationships between them, i.e. a undirected weighted correlation graph. The correlation plugin is automatically invoked when a .csv file is loaded and recognises attribute and data values as different data types. A user can transpose the matrix on import, so as to compare columns rather than rows, define the threshold for graph construction, scale or normalise data prior to calculation of the correlation matrix and add transforms prior to the visualisation of the resultant graph.

#### User Interface and Data Exploration

The Graphia user interface consists of an area displaying the graph itself, and a table listing the name and attributes of selected nodes. The graph display also shows the active transforms and visualisations, providing context to the graph displayed (Figure 2A). When using the correlation plugin, the table display includes a plot of the numerical data series associated with the selected nodes, upon which the correlation matrix was calculated (Figure 2B). The table can be adjusted such that only certain attributes are displayed, and the data plot area provides many options to modify the view, for example plot as line graph or histogram, individual node values or averaged, and various options to scale data, etc.. The graph display area and table/data plots areas can be viewed together or decoupled into separate windows. This enables display on separate screens where available.

#### Export of Results

Following or during an analysis it is possible to generate screenshots of the graph as displayed, lists of nodes and associated attribute data, data plots or analysis. Once an analysis session is complete the resulting graph can be saved in the tool’s own format (.graphia) which captures all the data used in its construction, and its layout, transforms and visualisations. A graph may also be exported to several other formats (pairwise text file, GraphML, GML), although in these cases information may be lost in the process.

#### Case Study 1: Visualisation of phylogenetic trees

Hierarchical data structures are often represented by tree graphs and used in biology to represent relationships between species, strains, samples or genes. While trees are an intuitive way of visualising such relationships, when the number of branches on the tree becomes very large, the ability to display such graphs at a local or global level is challenging. Here we show two examples of taxonomic trees visualised by Graphia representing the different levels of phylogeny, from a central node representing the class of organisms, up through branches representing the order, family, genus, with species and subspecies being the leaves of the tree. The examples described are taxonomic trees for all mammals and insects (Figure 3), as defined by NCBI Taxonomy database^23^. The taxonomic tree of mammals consists of 9843 nodes and 9862 edges and is shown in 2D with nodes coloured by type, i.e. what level of the taxonomic tree they represent. A small section associated with apes is highlighted (Figure 3B). When the graph is loaded using the WebSearch plugin, selection of a single node automatically initiates a search for the name of the selected node. Shown in Figure 3C is a taxonomic tree of all insect species, consisting of 275,328 nodes and 275,528 edges displayed in 3D. Graphia provides a unique environment to search, cluster and explore such data, the third dimension greatly assists in showing the structure of such large graphs.

**Figure 3.**
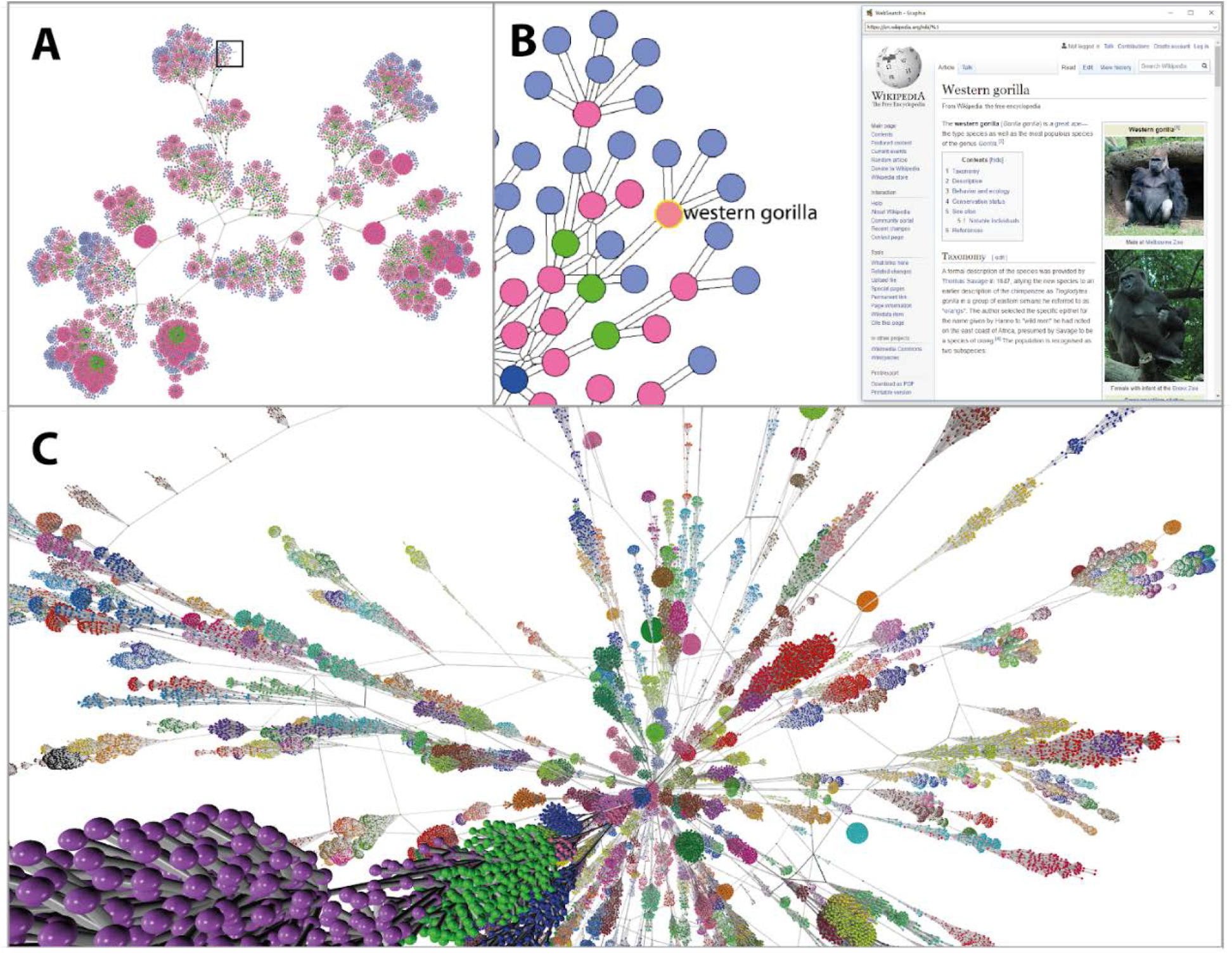
Visualisation of taxonomic trees. **(A)** Taxonomic tree of all mammals downloaded from the NCBI’s Taxonomy database, with nodes coloured according to type, i.e. subspecies (blue), species (pink), genus (green), etc. The graph comprises of 9,843 nodes and 9,862 edges and is shown with a 2D layout. **(B)** Zoomed-in view of the area in square shown in A, with a single node selected (Western gorilla) firing up a Graphia’s web-search plugin that automatically searches the web for the node’s name. **(C)** Taxonomic tree of all insects from the NCBI’s Taxonomy database, nodes coloured by Louvain cluster. The graph consists of 275,328 nodes and 275,528 edges and in this respect represents a large graph where visualisation in 2D is challenging.

#### Case Study 2: Analysis of single cell transcriptomics data

Single cell RNA sequencing (scRNA-Seq) generates gene expression profiles for thousands of individual cells in a single assay. The approach is an unbiased way of identifying cell types in a mixed population of cells, both known and uncharacterised, in addition to the genes that define them. This has seen a surge of interest in dimensionality reduction methods, in particular t-SNE^24^ and UMAP^25^, as users seek to optimise the visualisation of results (Figure 4A,B). These methods are constrained by issues associated with representing the underlying data structure, i.e. relationships between cell groupings and when 10s of thousands of cells are analysed, a 2D plot space is limiting.

**Figure 4.**
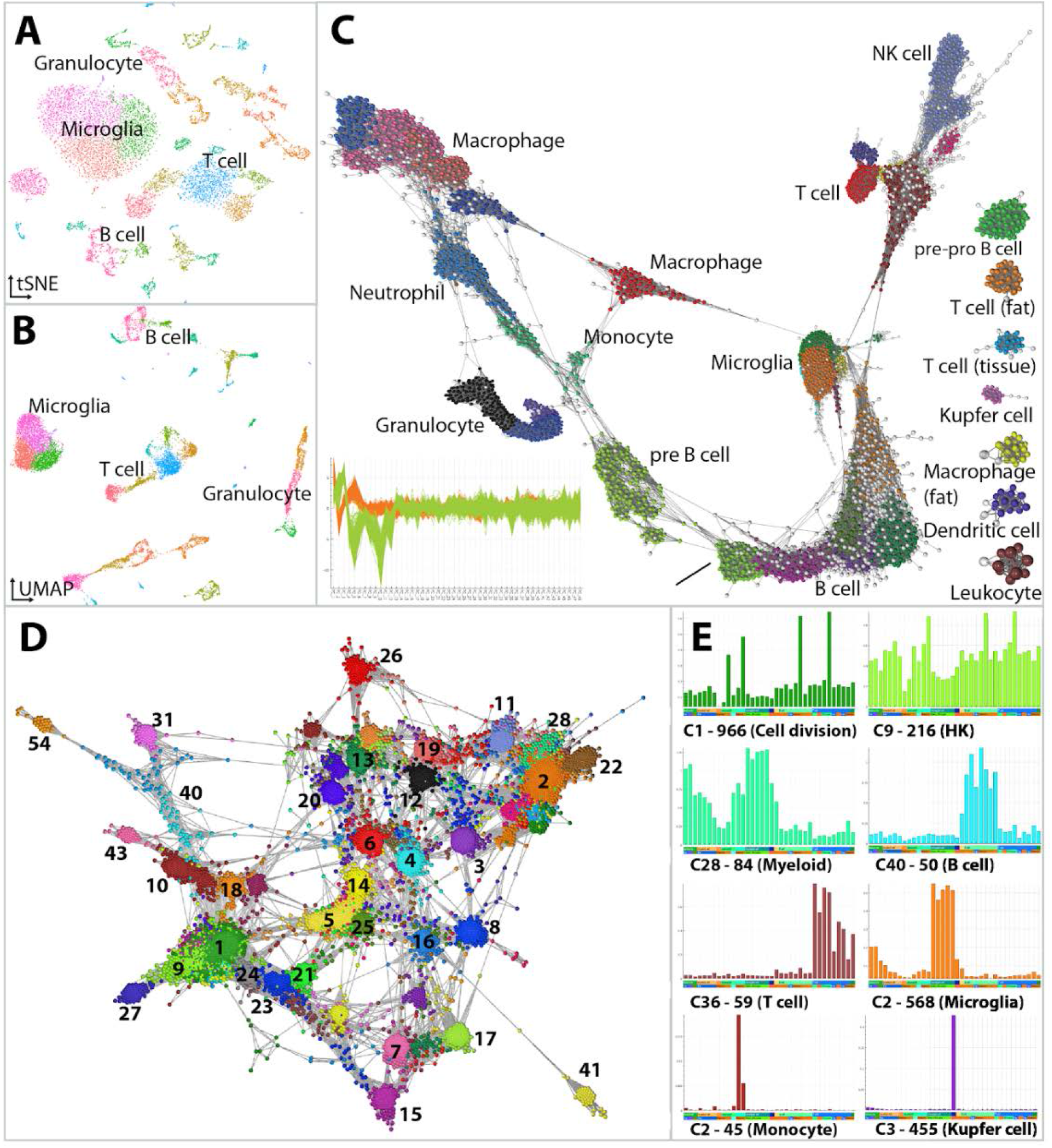
Analysis of cell and gene associations in scRNA-Seq data. The structure of scRNA-Seq data is commonly represented using approaches such as **(A)** t-SNE and **(B)** UMAP as shown here for immune cells derived from the Tabula Muris dataset. However, the distance between data points and groups of data points is difficult to interpret. **(C)** Graphia enables the construction of cell-to-cell networks built on a similarity parameter. Here, the 48 most significant PCA values for each cell were first calculated and this PCA profile used to construct a correlation network. The plot bottom left of C, shows the PCA profiles of cells in the two largest cell clusters. To better show graph structure, a k-NN (k = 10) was applied and outlier cells removed (*r* < 0.85 and node degree < 10, nodes coloured white). The graph comprises of 12,498 nodes (cells) and 143k edges. Cell clusters have been annotated as the cell types defined by the authors. **(D)** Shows a gene correlation network generated from these data by first calculating the average expression of genes within cell clusters and then calculating a correlation matrix from these values. **(E)** Plots show the average expression profile (y-axis) of a selection of gene-clusters across the aggregated cell-clusters (x-axis). The label gives the cluster number, e.g. C1, the number of genes within the cluster (- 966) and the association of the genes with a given biology or cell type.

An alternative approach is to treat single cell data as a graph, where nodes represent cells or genes, and edges the similarity between them. There are a number of measures that may be used to calculate the distance between cells or genes and here we discuss our currently favoured approach. The Tabula Muris dataset^26^ includes a scRNA-Seq data from 20 different mouse tissues. For the purpose of this study we selected only data from tissue immune cells, as annotated by the authors. Preprocessing and quality control was performed as per the Tabula Muris pipeline (https://github.com/czbiohub/tabula-muris), producing a normalized dataset of 14,466 cells from 12 tissues. Principal component analysis (PCA) was conducted to reduce the gene profile of a cell to principal components (PC), with the 48 most significant PCs (*adj. P-value* <0.05) based on Jackstraw permutations^27^ being considered. This file (cells as rows, PCs as columns) was loaded into Graphia and a graph was generated based on the Pearson correlation between the PC profile of cells. An initial network was generated applying the k-NN algorithm (k = 15) and outlier cells were removed. Outliers were defined as cells with poor correlations and connectivity with other cells of the same cell type (*r* < 0.85 and node degree < 10). Applying these filters produced a network that better separated known cell-types. Subsequently, the filtered 12,498 cell-to-cell network was clustered using the Louvain clustering algorithm^22^ with a granularity of 0.8, identifying 36 cell clusters (Figure 4C).

Following the identification of cell clusters, it is generally of interest to identify gene markers and expression modules associated with different cell types. Due to the inherent noise in scRNA-Seq data, it is of limited use to construct gene networks based on the correlation between expression values of individual cells. Instead, one alternative is to construct gene correlation networks based on an aggregated gene expression value across cells from each cluster, i.e. the similarity between clusters of cells not individual cells. Accordingly, a matrix of the average gene expression values across clusters defined above was loaded into Graphia. A network of strongly correlated genes (*r* > 0.85) was generated and gene coexpression modules identified using the Markov clustering algorithm^21^ with a granularity setting of 1.7 (Figure 4D). Graphia enables a dynamic and rapid exploration of these clusters, allowing a user to understand where a given cluster sits within the context of the entire graph, the identity of genes present within a given cluster and the profile of all or some of those genes across samples, in this case cell clusters (Figure 4E).

#### Case Study 3: Exploration of bacterial pangenome structure

Whole genome sequencing is now routine in many fields. One common use is in the characterisation of microbial species and public databases which already hold tens of thousands of genome sequences for the best studied organisms. Graph-based methods and tools that support the visualisation and analysis of such data are well established. For instance, Bandage is a software package now widely used to visualise *de novo* assembly graphs of bacterial genomes^28^. PPanGGOLiN^29^ and Panaroo^30^ are graph-based pangenome clustering tools for the analysis of genomic diversity within a bacterial species (i.e. its pangenome), which can then be used to statistically classify genes according to their occurrence in the genomes.

Comparative analyses of bacterial sequences have revealed a high degree of genetic diversity between isolates of the same organism, leading to the concept of “core” genes present in all isolates and “accessory” genes present only in some isolates. The distribution and organisation of accessory genes has a significant impact on an organism’s ability to adapt to different hosts and niches, virulence and drugs. Figure 5 shows a graph generated from a previously published dataset of whole genome sequencing data of 778 *Staphylococcus aureus* isolates^31^. Genomes were annotated using Prokka (v1.13)^32^ and their pangenome defined using the analysis pipeline PIRATE (v1.0.3, default settings used, 95% sequence similarity threshold)^33^. In this visualisation, nodes represent individual genes/gene variants and edges the syntenic relationships between them. When the visualisation is enhanced through making node/edge size and colour proportional to the number of genomes in which a gene is present or two genes are syntenic across the dataset, core regions of the genome become easy to identify (Figure 5A). These can also be collapsed to single nodes to simplify the graph. Similarly, areas of high variability within the pangenome are obvious (Figure 5B). Graphia can be used to identify specific nodes, for example the path of a single genome (RF122) can be shown in the context of the wider pangenome (Figure 5C) or used to explore smaller local variations, e.g. the quorum sensing locus *agrABCD*, which shows four variants when defined at this similarity threshold (Figure 5D,E). In principle, displaying variation between sequences as a network is applicable to any such data. This has relevance to “pan-reference” genomes for more complex species such as humans, as well as for showing other variation, such as clustering repeat regions^34^ or alternative isoforms of transcripts^35^.

**Figure 5.**
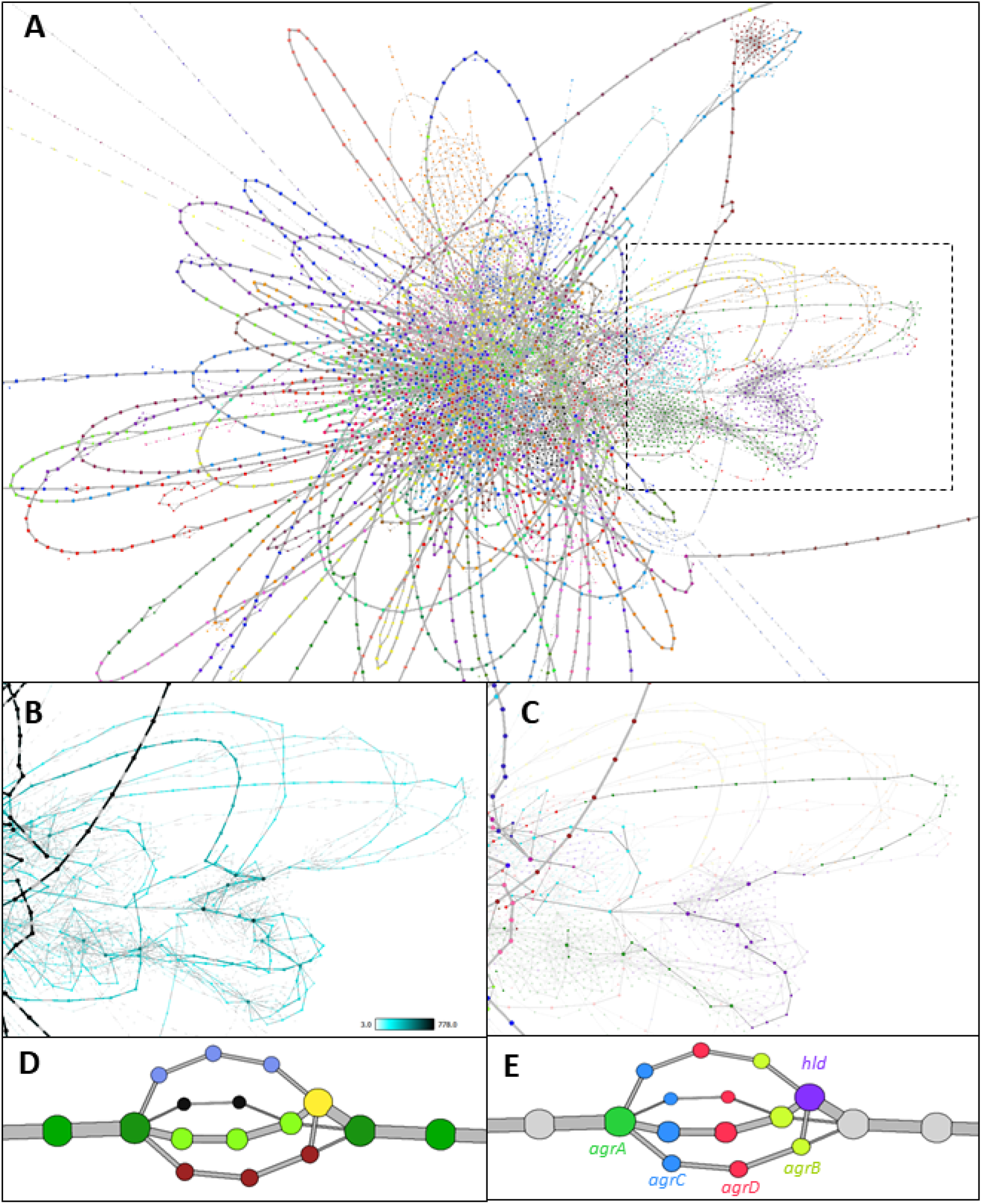
Visualisation of the pangenome of *Staphylococcus aureus*. **(A)** The full pangenome of 778 isolates. Nodes represent individual orthologous genes as identified by PIRATE. Node size is determined by the number of genomes in which a gene has been identified. Edges denote where two genes are syntenic, and their thickness is determined by the number of times this syntenic connection is observed across isolates. Syntenic stretches of core genes have been collapsed for clarity using the “Contract Edges” transform, and low confidence nodes and edges (*n* < 3) have been removed. Coloured by Weighted Louvain Clustering (granularity = 1.0). **(B)** Highly variable region (boxed area in A) with a high density of “phage-like” genes. Nodes and edges are sized and coloured by frequency. **(C)** Genes highlighted are all found in single *S. aureus* isolate, RF122. **(D)** the *agrABCD* locus coloured by gene-association clustering. Frequently, an alternative allele is not identified as being the same gene, but their position is strongly indicative of shared function. **(E)** the same locus coloured by gene identity.

#### Case Study 4: Analysis of Human Genome Variation

Genome variation also occurs at the level of individual DNA base pairs, so called single nucleotide variants (SNVs). Genotypes can be scored in an individual in terms of their allele dosage, i.e. both being the same as the reference nucleotide (0), heterozygous (1) or homozygous for the variant (2). Calculation of the correlation across a range of these positions in a population of individuals creates a relationship matrix.

Figure 6 shows various views of data from the 1000 genomes project^36^. Here 23,675 SNVs from chromosome 22 were used to generate graphs of the relationships between individuals in this cohort and the SNVs themselves. These data represent the genomes of 2,504 individuals, who were selected 26 distinct ethnic populations from five continents. The average correlation between the SNV profile of individuals is low and the graph shown in Figure 6A is constructed using a threshold of *r* = 0.238, a value at which most of the genomes in the cohort form one connected component. To reduce the edge count (from 400.4k to 5996) and open-up the local structure the k-NN algorithm was applied such that only the top three relationships between individuals were maintained. The topology of the graph is clearly strongly influenced by the ethnicity of individuals with discrete clusters being observed for all the five continental populations and in some cases individuals from certain countries or ethnicities showing a local grouping within this overall structure. Also visible from the graph are a number of very strongly related individuals (Figure 6Ai) and a number of instances where an individual does not co-occur with their annotated population, e.g. there are a number of South Americans in with the Africans, and vice versa (Figure 6B). Transposing the matrix to analyse the similarity between the profile of SNVs across the 2,504 individuals, at the threshold used here (*r* = 0.75), 11,600 SNVs formed 2,467 separate graph components of more than one node (Figure 6C). After clustering the graph using the Louvain algorithm, many of the clusters contain nearby SNVs; likely haplotype blocks, some of which were clearly associated with a given a population. Graph analysis represents an improved approach compared to e.g. PCA plots, visualising genetic associations between individuals and genetic variants.

**Figure 6.**
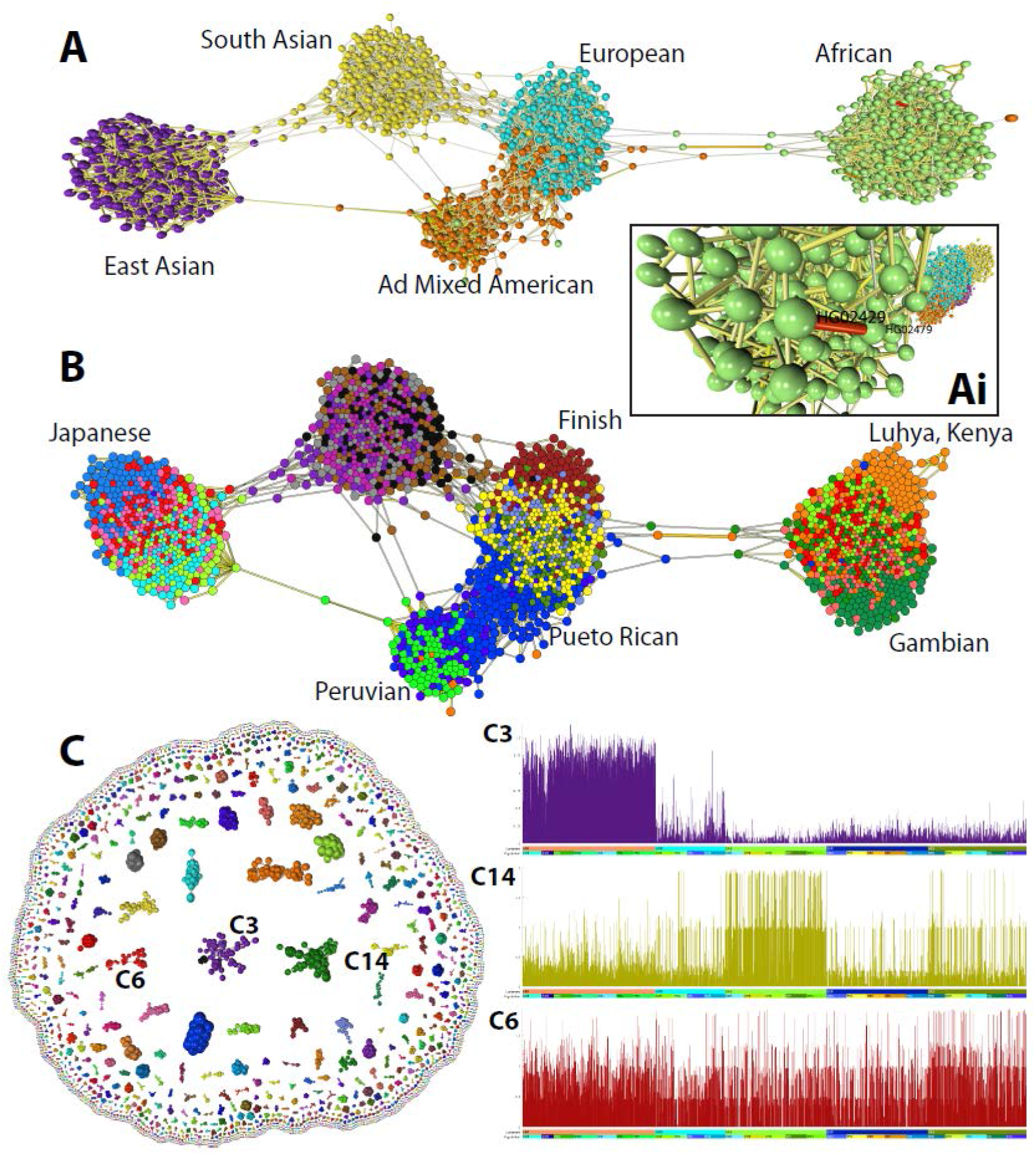
Analysis of Single Nucleotide Genome Variant Data. **(A)** The graph shown was constructed from data from the 1000 genomes project based on the correlation (*r* threshold = 0.238) between the allele dosages at 23,675 SNVs from chromosome 22. Nodes represent the 2,504 individuals included in the study and edges the three most significant correlations with their neighbours (k-NN was applied where k = 3). In most cases, individuals’ group with others from the same continent although there are instances where this does not appear to be the case. Visualisation of edge weights **(Ai)** also highlights cases where individuals would appear to be closely related. **(B)** Colouring of nodes by the attribute ‘population’ provides a higher resolution to the graph and populations showing a high degree homogeneity have been labelled. **(C)** Transposing the data upon import demonstrates SNVs whose pattern across the genome covaries. Clustering of these data shows many to represent haplotype blocks and inspection of their profile across genomes, demonstrates some SNV clusters to be associated with a given ethnic grouping, e.g. cluster 3 (Africans) and cluster 14 (East Asia), whilst others little obvious association with ethnicity, e.g. cluster 6. Plots show the average score of SNV’s within a cluster (y-axis, 0,1,2), across the 2,504 individuals ordered by continent and then population (x-axis).

## DISCUSSION

Data-driven research is now a foundation of modern biomedical and agricultural sciences, due to continued growth in the size and complexity of biological datasets. Network analysis provides a flexible toolbox combining visualisation with the algorithmic analysis of data structure, for testing a broad range of hypotheses and hypothesis-free data explorations. However, when datasets become very large existing network tools struggle both to render the number of graph elements (nodes/edges) and to arrange them in a navigable format for human interaction. Graphia is designed for the visualisation and analysis of large graphs. Originally, our interest in graph-based analysis was driven by our desire to analyse large correlation networks of transcriptomics data. The results of weighted gene coexpression analysis (WGCNA)^3^ are generally visualised as a tree diagram or heat-map. The precursor of Graphia, BioLayout *Express*^3D 4,6^, was developed specifically to generate and display transcriptomic data and pathway modelling^4,37,38^. BioLayout has been used in the analysis of many large transcriptomic datasets from multiple species^39–43^. It has also been applied to datasets that were not envisaged at the time, for example the relationship between symptoms of altitude sickness^44^, the honey bee microbiome^45^, comparing morphometric measurements of dog brains^46^ and even naming patterns in historical birth records^47^. The addition of new functionality to BioLayout was however constrained by inherent limitations in the code structure and programming language (Java).

Graphia is an entirely new platform developed using a modern UI framework (Qt) and programming language (C++). The correlation plugin reproduces and improves upon the functionality of Biolayout for the analysis of any high dimensional numerical matrix. Data visualisation is core to the functionality of Graphia as an analysis platform. Good visualisations make it easier for a user to recognise patterns, trends, and outlier groups within data. The next step in an analysis is determined by insights gained from the interaction with the visualisation, whether that be the discovery of errors in the input data, data effects due to technical reasons, or from new and interesting discoveries. Graphia is designed to make best use of the latest accelerated graphics hardware, to make graph visualisations scalable, but still responsive in real time. By default, graphs are rendered in 3D, where the visualisation and navigation of complex graph topologies is much enhanced; the additional dimension providing the ability to distinguish the distance between what might appear in 2D to be closely connected nodes. Another core aspect to the visualisation of data is the concept of graphs being ‘dynamic’, changing in real-time, as nodes/edges are added or removed. To achieve this, the layout algorithm runs continuously, unless manually paused. Dynamic transitions may become challenging when graph structure alters dramatically following a transformation, such as when a hub-node is deleted from a tree graph or one graph component fragments into many. If such a transformation is executed quickly, a user’s ‘mental map’ can be lost^48^. For this reason, Graphia includes the option to slow down the transition between one state and the next, and in addition orientates components ‘in flight’ prior to their reconnection. Indeed, the way in which Graphia handles graph components dynamically is quite unique, from their concentric layout, to the fact that even singletons can be rendered.

The development of Graphia has been driven by the analytical challenges associated with data derived from the biological sciences, but it is designed as a general-purpose platform for the analysis of network data from any source. If the input data is a table of numerical values (continuous or categorical), a distance matrix can be used to build a graph, or if data already exists in a graph format, Graphia provides a means to explore it. Graphia currently loads data from files, but in theory it could be loaded from a web resource or local database. It is interesting to note the widespread adoption of graph databases. Not only do graph databases speed up and simplify querying of data stores, storage as a graph makes it easier to visualise and analyse it. Whilst there are a growing number of web-based tools that support the querying and visualisation of graph databases, none possess the power of Graphia in rendering large portions of the data they store.

Here we offer a high-level view of the functionality of Graphia and some examples of its many uses within the biomedical sciences. We provide installers to allow it to run on all common desktop operating systems and the access to the source code, to allow users to develop new functionality to enhance its functionality for their needs.

## Contributions

TCF, TA and SH designed Graphia, TA and SH wrote the code. AP, JH-L, TR, BS, DAH and JP developed the use cases shown here. All authors contributed to the testing of the software and writing of the paper. User support was developed primarily by TCF, TA and SH.

## Acknowledgements

Graphia was originally designed and built by Kajeka Ltd., a University of Edinburgh spin-out company (2015-2020). We would like to acknowledge all those who supported this venture, in particular grant funding from Scottish Enterprise (SMART/14/034 / 14/9168). During the period of Graphia’s development TCF and JP were supported by the Roslin Institute’s Strategic Grant from the UK’s Biotechnology and Biological Sciences Research Council (BBSRC) [BBS/E/D/10002071].

